# Nonlinear spatial integration allows the retina to detect the sign of defocus in natural scenes

**DOI:** 10.1101/2024.06.03.596421

**Authors:** Sarah Goethals, Awen Louboutin, Samy Hamlaoui, Tom Quetu, Samuele Virgili, Matias A. Goldin, Konogan Baranton, Olivier Marre

**Affiliations:** Groupe Lens Innovation, R&D Life and Light Science, EssilorLuxottica, Paris; Institut de la Vision, Sorbonne Université, INSERM, CNRS, Paris; Groupe Lens Innovation, Digital Innovation, EssilorLuxottica, Paris

## Abstract

Eye growth is regulated by the visual input. Many studies suggest that the retina can detect if a visual image is focused in front or behind the back of the eye, and modulate eye growth to bring it back to focus. How can the retina distinguish between these two types of defocus? Here we simulated how eye optics transform natural images and recorded how the isolated retina responds to different types of simulated defocus. We found that some ganglion cell types could distinguish between an image focussed in front or behind the retina, by estimating spatial contrast. Aberrations in the eye optics made spatial contrast, but not luminance, a reliable cue to distinguish these two types of defocus. Our results suggest a mechanism for how the retina can estimate the sign of defocus and provide an explanation for several results aiming at mitigating strong myopia by slowing down eye growth.

## Introduction

In most animal species during development, the eyes grow alongside with the body. The eye growth process is highly constrained since it should maintain sharp vision all along development. This means that the eye needs to constantly adjust its axial length to the optical power of the eye optics, to ensure proper image focus on the retina, a process called emmetropization.

The visual input is the main regulator of this homeostatic process (1). Previous works have shown that in many species during development, eye growth can be perturbed by modifying the visual input (reviewed in (2)). In young animals, placing a positive lens in front of an emmetropic eye focuses the image in front of the retina and it was observed that, to compensate for the imposed defocus, the eye slows down its growth rate (originally shown in chicks in (3), then also in rhesus monkeys (4), marmosets (5), tree shrews (6), guinea pigs (7) and mice (8, 9)). On the other hand, placing a negative lens in front of an emmetropic eye focuses the image behind the retina. In this situation, to compensate for the imposed defocus, the eye accelerates its growth rate (shown in chicks (3, 10), rhesus monkeys (11), marmosets (12, 5), tree shrews (13, 14), guinea pigs (7) and mice (8)). These studies show that, during development, the eye is able to compensate for an imposed defocus to become emmetropic. Furthermore, a large body of work has shown that excessive eye elongation can also be induced by strongly degrading the visual input (reviewed in (2)).

How does the eye adapt its growth rate to become emmetropic? Several studies strongly suggest that the retina plays a central role in controlling eye growth and that feedback from the brain is not necessary. Sectioning the optic nerve in chicks does not prevent the acceleration of growth rate in response to an imposed positive defocus (15), nor does it prevent excessive eye growth in response to visual input degradation (16–19). Moreover, blocking action potentials from retinal ganglion cells (RGCs) by intravitreal injection of tetrodotoxin (TTX) does not prevent eye elongation in chicks (20) and tree shrews (21) that underwent visual input degradation. Finally, depriving only half of the visual field results in local eye elongation in the corresponding half of the eyeball (22, 23), and eye growth seems to be mostly driven by the peripheral retina (24–26). All together, these results suggest that eye elongation is driven by retinal mechanisms that operate locally (27).

This means that the retina is able to distinguish an image focused in front of the retina from an image focused behind the retina, in order to trigger the appropriate change in growth rate when a defocus is imposed. How the retina achieves this is not clear.

Here we developed a new hybrid approach combining optics, electrophysiology and modelling to investigate how the eye optics influence mice retinal cell’s activity. On the optics side, we simulate the optics of the mouse eye with a detailed optical model, for different positions of the focal point with respect to the retina. Then we transform natural images with these optics, to obtain realistic defocused retinal images. On the electrophysiological side, we dissected the retina to remove the real optics of the eye, projected the defocused images on the *ex vivo* retina and recorded the impact of the sign of defocus on the retinal ganglion cell’s activity. We identified two types of RGCs whose activity systematically decreases with a more positive defocus, independently of the specific visual input. This shows that some RGCs can robustly detect the difference between an image focused in front and behind the retina. By modeling the responses of these cells, we found that they compute local spatial contrast (28) inside their receptive field, and we show that computing local spatial contrast is sufficient to detect the sign of defocus. This property arises from the spherical aberrations of the eye optics that make this contrast very different for positive and negative defocus. Our work suggests a strategy for the retina to determine the sign of defocus and provides an explanatory basis for several results in the literature concerning myopia mitigation strategies that aim at slowing down the eye growth.

## Results

### Simulating the mouse eye optics

To measure the impact of the eye optics on retinal activity, we designed a hybrid approach: we simulated in details how natural images are transformed by the optics of the eye. We then projected these transformed images on the retina *in vitro* and measured how ganglion cell’s responses were influenced by the optical transformation.

The experiments were done in retinas of mice, an animal model in which eye growth can be perturbed (8, 9), ganglion cell activity can be recorded (29), and the eye optics has been characterized (30, 31).

Our detailed model of the mouse eye optics (see Methods) takes into account optical aberrations. As a result, and in contrast with a model with no aberrations, the image projected on the retina can be different if the image is focused in front of the retina or behind. To illustrate this, we simulated how a point source of light transformed by the eye optics appears on the retina, which is usually called the Point Spread Function (PSF) of the optical system. It represents a local linear approximation of the transformation imposed on the stimulus by the eye optics. This PSF will depend on the position of the focal point with respect to the retina, and on the eccentricity (in degrees of visual angle) of the spot of light. Note that, in our model, the eye optics is fixed and the defocus is simulated by moving the position of the retina with respect to the focal point (fig. 1A). For various eccentricities, the PSF of an image focused in front of the retina was always different from the PSF corresponding to an image focused behind the retina (fig. 1B). The PSF spread around the main focal point is much broader for a focus in front of the retina. For simulated peripheral eye optics, the PSF spreads around the main focal point mainly in one direction along the horizontal axis (fig. 1B). When convolving these PSFs with natural images, the change of focus had a visible impact on the resulting image falling on the photoreceptor plane: the contrast and the amount of blur were different if the focus was positive (focal plane in front of the retina) or negative (focal plane behind the retina).

**Figure 1:**
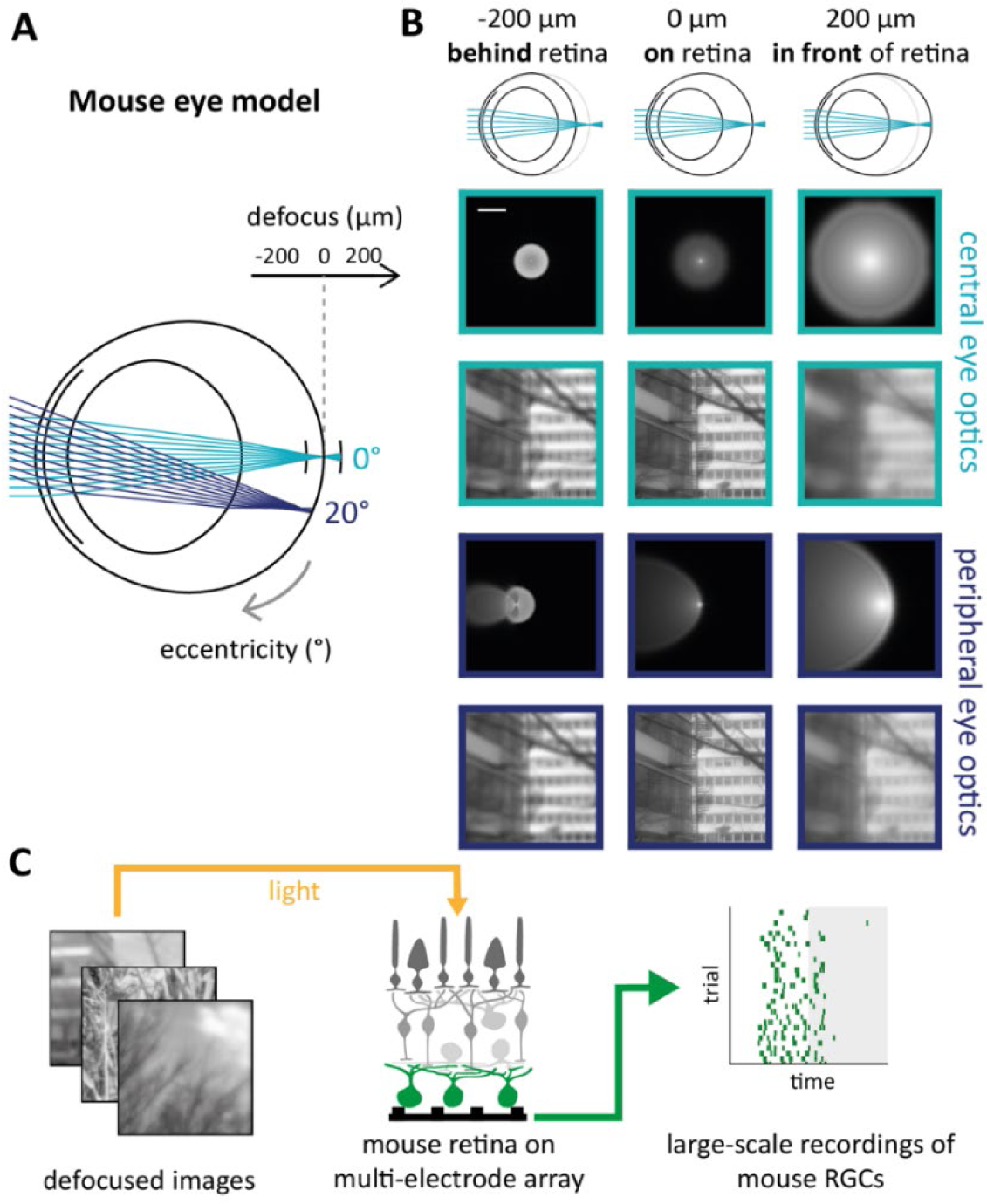
A mouse eye model to simulate retinal images. **A**, the mouse eye model and its parameters. First, the light source eccentricity: light rays can reach the retina in the centre (0°, light blue) or periphery (20°, dark blue). Second, the position of the retina with respect to the focal point (defocus, in µm). Indeed, in the model, we kept the eye optics fixed and simulated a change of defocus by moving the position of the retina. Negative values of defocus correspond to the image being focused behind the retina, and positive values of defocus correspond to the image being focused in front of the retina. **B** Top, mouse eye model when the image is focused behind (left), on (centre) or in front (right) of the retina. Bottom, Point Spread Functions (first and third rows) and the corresponding retinal image (second and fourth rows) for simulated central (0°, light blue) and peripheral (20°, dark blue) eye optics, when the image is focused behind (left), on (centre) or in front of the retina (right). **C**, experimental preparation. Natural images transformed by the eye optics (left) are projected on an *ex vivo* mouse retinal flattened on a multi-electrode array (centre), to record RGC’s responses (right). RGCs are represented in green. The raster plot on the right is the spiking activity of one cell in response to 30 presentations of the same image during 300 ms. The grey area corresponds to the presentation of a grey frame during 300 ms. Credit for the natural images shown here goes to Hans Van Hateren (67).

The images transformed by the eye optics will thus be different if the image is focused in front or behind the retina. Note however that the PSFs were shown in log-scale, which amplifies these differences. It is thus unclear if the retina can detect this difference, in particular when stimulated with natural images.

### Some types of mice RGCs detect the sign of defocus

To answer this question, we projected these defocused images on mice retinas *ex vivo* and recorded RGC’s electrical responses to different optical transformations with a multi-electrode array (MEA) (fig. 1C, see Methods). We then measured how RGC’s firing rate varies with the simulated defocus, simulating different eccentricities to test the robustness of our results. In addition, we presented a chirp stimulus (fig. 2A) to classify RGCs depending on their response to full-field bright and dark stimulus (see Methods) (fig. 2B).

**Figure 2:**
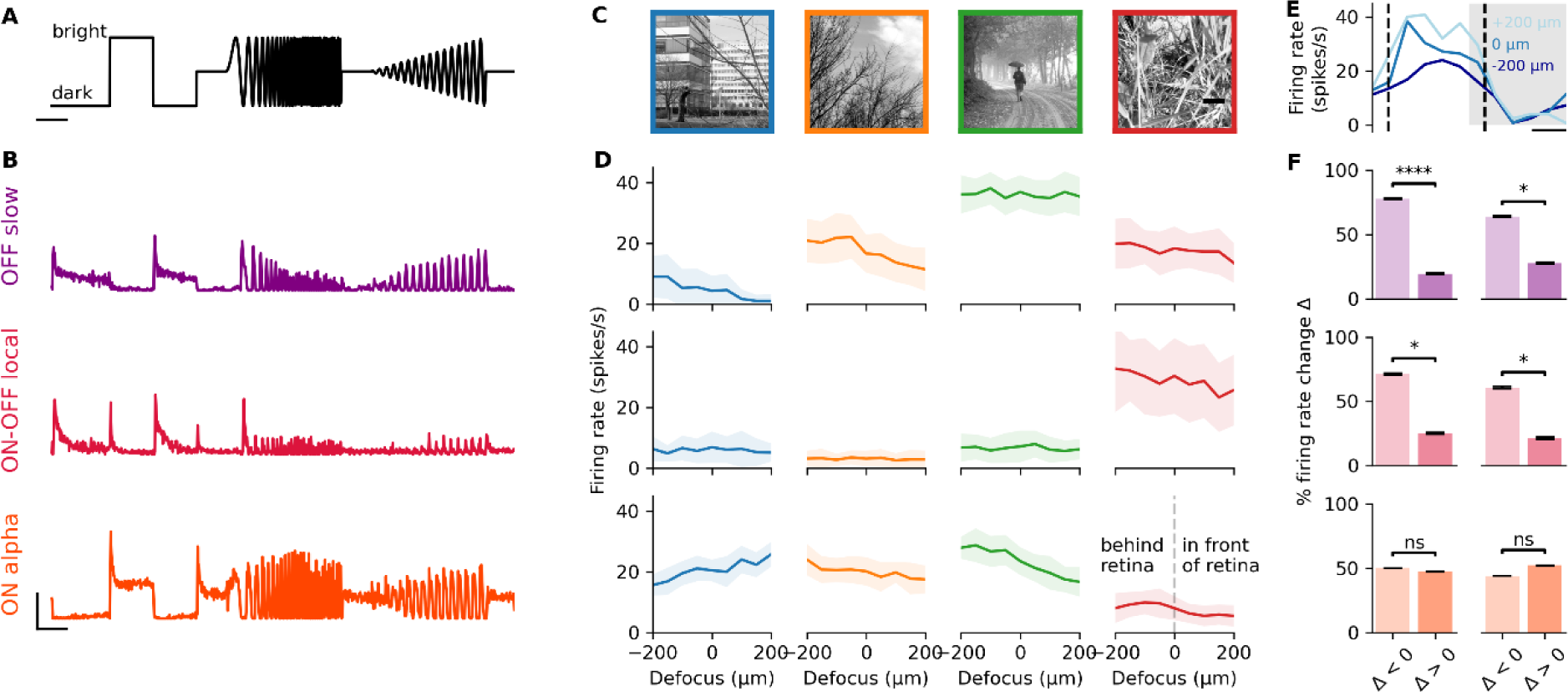
Specific types of retina ganglion cells could signal the sign of defocus. **A**, chirp stimulus (see Methods). Scale bar: 2 seconds. **B**, example RGC response to the chirp stimulus. Top, OFF slow; middle, ON-OFF local; bottom, ON alpha. Horizontal scale bar: 2 seconds. Vertical scale bar: 20 spikes/second. **C**, the 4 natural images that were defocused and flashed on the mouse retina during MEA experiments. Scale bar: 500 µm. **D**, firing rate as a function of defocus for the images in panel C and for three example cells, for simulated peripheral eye optics (20°). Top, OFF slow; middle, ON-OFF local; bottom, ON alpha. Data represented as mean ± standard deviation. **E**, Example of the raw response of an ON alpha cell to the “blue” image: firing rate during image presentation (white area) and grey frame presentation (grey area) for a defocus of –200 µm (dark blue), 0 µm (blue) and 200 µm (light blue). The vertical dashed lines represent the time range during which we measure the firing rate. Note that that cell’s firing rates increases with defocus, as the ON alpha example cell from panel D. **F**, proportion of images and cells leading to a firing rate change Δ (between defocus of +200 µm and defocus of –200 µm), that is strictly negative (left bar plot) or strictly positive (right bar plot). Left column: simulated central eye optics (0°). Right column: simulated peripheral eye optics (20°). Top, OFF slow (N = 9 cells x 4 images), centre: p = 5 × 10^-5^; periphery: p = 1 × 10^-2^. Middle, ON-OFF local (N = 7 cells x 4 images), centre: p = 1 × 10^-2^; periphery: p = 1 × 10^-2^. Bottom, ON alpha (N = 55 cells x 4 images), centre: p = 0.1; periphery: p = 0.9). One-sided Wilcoxon signed-rank test. Credit for the natural images shown here goes to Hans Van Hateren (67).

We found that for many cells, the change in the simulated defocus was often enough to evoke a change in their firing rate (fig 2C, D, E). The difference in firing rate was visible when switching from a positive defocus (focus in front of the retina) to a negative defocus (focus behind the retina) (fig. 2E). The changes in image properties induced by the change in PSF are thus sufficient to evoke a change in the RGC response.

Presumably, the sign of this firing rate difference should depend on the content of the image inside the receptive field of the cells, and this is indeed what we found for many of them. However, surprisingly, we found two RGC types (OFF slow and ON-OFF local (32)) (fig. 2B) where the firing rate almost never increased when the defocus changed from negative to positive: it always decreased or stayed constant, for the images we tested (fig. 2C, D), across all simulated eccentricities of the optical model, and across all the cells of the same type (fig. 2F) (OFF slow, N = 9 cells x 4 images, centre: 78 vs. 19%, p = 5 × 10^-5^, periphery: 64 vs 28 %, p = 1 × 10^-2^; ON-OFF local, N = 7 cells x 4 images, centre: 71 vs 25%, p = 1 × 10^-2^, periphery: 61 vs 21%, p = 1 × 10^-2^, one-sided Wilcoxon signed-rank test). This suggests that these cells could reliably detect the sign of defocus.

Note that we found this behaviour in RGCs from other types (see Discussion), but this was not the case for all the cell types. ON alpha cells, for example, did not show this property; their firing rate could either increase or decrease when the sign of defocus changed, depending on the natural image displayed (fig. 2F, ON alpha: N = 55 cells x 4 images). Therefore, the systematic decrease property is specific to some cell types and not shared by all RGCs.

We have found that, for these two types of RGCs, firing systematically decreases with defocus. To test if this decrease is systematic over a very large set of images, we built a model to predict how these RGCs may respond to other defocused natural images. We used a convolutional neural network (CNN) model of the retina that has been previously shown to predict well ganglion cell responses to natural images (33) (fig. 3A). The model was trained on the RGC’s responses to a large set of 3010 not defocused natural images (see Methods). Its performance was evaluated on a set of 30 held-out repeated sharp natural images (‘test set’). For both types this CNN model accurately predicts RGC’s response to sharp natural images (OFF slow: R² = 0.76 ± 0.13; ON-OFF local: R² = 0.81 ± 0.13; ON alpha: R² = 0.96 ± 0.02) (fig. 3C). It could also generalize and predict with comparable performance RGC’s response to defocused images (OFF slow: R² = 0.92 ± 0.06; ON-OFF local: R² = 0.77 ± 0.3; ON alpha: R² = 0.97 ± 0.02) (fig. 3C).

**Figure 3:**
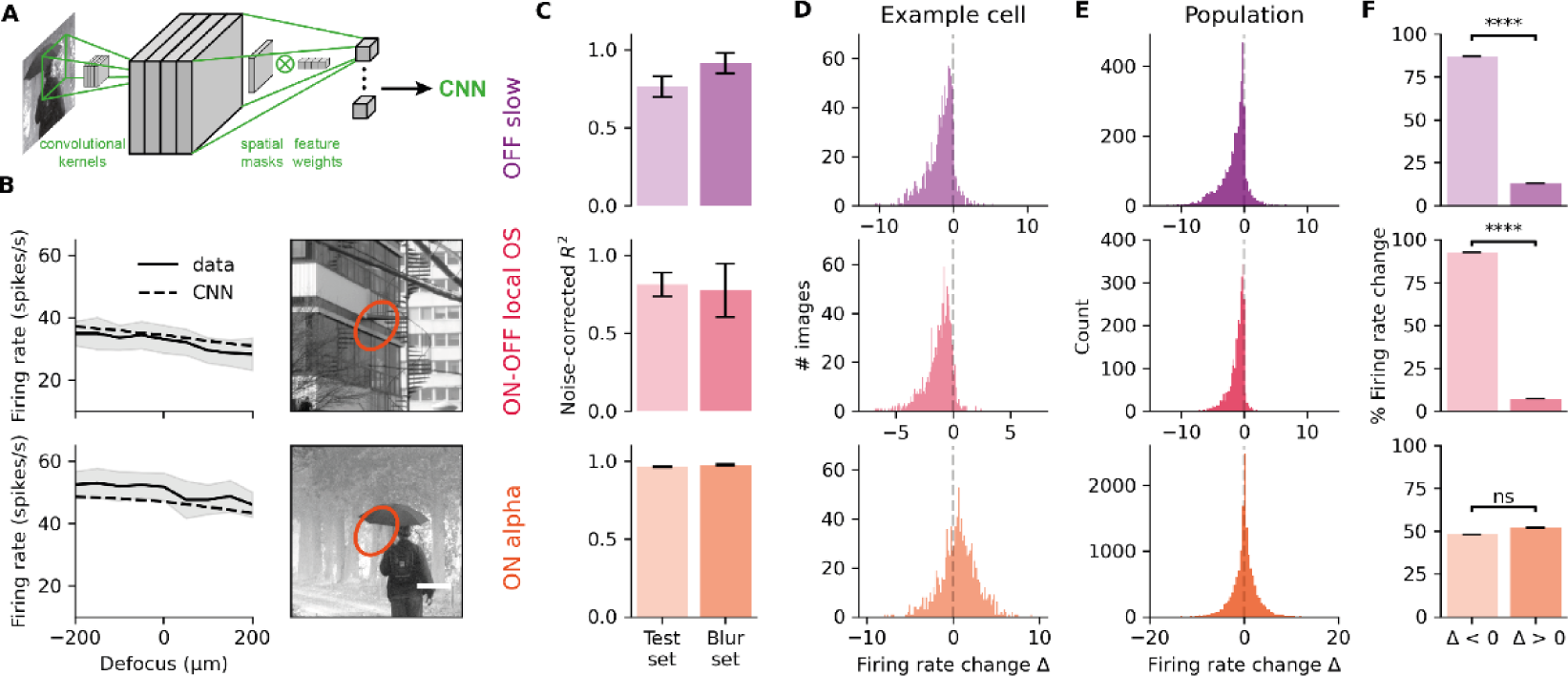
A convolutional neural networks model accurately predicts RGCs response to defocused images and confirm that two types almost always decrease their activity when the focus switched from negative to positive. **A**, Schematic of the CNN model architecture. Adapted from (Goldin et al., 2022). **B**, Example of an ON alpha cell’s response (full lines) to the images shown on the right for simulated peripheral eye optics (20°), and the predictions of the CNN model (dashed lines). The orange ellipse on the images is the receptive field of the cell. Scale bar, 200 µm. **C**, Average performance of the model at predicting the response to repeated sharp natural images (left) and repeated defocused images (right). Data from N = 4 OFF slow cells (top), N = 3 ON-OFF local cells (middle) and N = 15 ON alpha cells (bottom). Data are represented as mean ± SEM. **D**, distribution over N = 1000 images of the change in predicted firing rate Δ between defocus of 200 and –200 µm for an OFF slow (top), an ON-OFF local (middle) and an ON alpha (bottom) example cells. **E**, same as D for all the modelled cells. Top, OFF slow (N = 4 cells x 1000 images). Middle, ON-OFF local (N = 3 cells x 1000 images). Bottom, ON alpha (N = 15 cells x 1000 images). **F**, proportion of images and cells leading to a firing rate change Δ (between defocus of +200 µm and defocus of –200 µm), that is strictly negative (left bar plot) or strictly positive (right bar plot). N is the same as for panel D. Top, OFF slow (p = 0); middle, ON-OFF local (p = 0); bottom, ON alpha (p = 0.1); one-sided Wilcoxon signed-rank test. Data of **D, E** and **F** are shown for the simulated peripheral eye optics (eccentricity = 20°).

We then used this model to test whether the cell’s firing rate varies in the same direction whatever the specific image. We predicted RGC’s responses to thousand defocused natural images and examined the distribution of the change in firing rate between a positive defocus and a negative defocus (Δ) (fig. 3D, E). For the OFF slow type, 87.1% of the images across the cell population lead to a negative change in firing rate, and this proportion is 92.7% for ON-OFF local cells (fig. 3F). It shows that that these cell’s firing rate is strictly lower for a positive defocus than for a negative defocus, for almost all natural images. On the contrary, ON alpha cell’s firing rate did not show this effect: when the defocus changed from positive to negative, firing rate could either decrease or increase depending on the natural image, with equal probabilities (48.0% vs. 51.2%) (fig. 3F, see also the almost symmetrical distribution of the change in firing rate fig. 3E).

To summarize, the CNN model predictions confirm that some types of RGCs almost always decrease their firing rate with defocus, whatever the specific visual input, and that this is a specific property of some type, but not ubiquitous among ganglion cells. Our results suggest that these cell types can be used to robustly detect the sign of defocus during natural image stimulation. From now on we will call them « defocus detectors ».

### Defocus detector cells encode local spatial contrast

We next wondered how can defocus detectors actually perform the detection of the sign of defocus. Presumably, they have to extract a specific feature from the natural image, that will unambiguously tell the sign of defocus. To answer this question, we extracted the values of the mean intensity and contrast inside the receptive fields (most studied low level image statistics) of the two cell types identified above, across different images and eye model parameters, and used them as inputs to simple models to study if they could reproduce our previous results.

We first learned a classical linear non-linear model (28), which takes as an input the mean intensity inside the receptive field centre and aims at predicting the response (“Intensity model”, fig. 4A). The parameters of this model were learned from the response of each cell to a set of flashed natural images (same set as for the CNN, see Methods). Following Liu and colleagues (28), we then tested whether including second order statistics as an input to the model could improve the performance (fig. 4B). For each image presented, we measured the local spatial contrast (LSC, see Methods), which is the standard deviation of the pixel intensity in the receptive field centre. When this spatial contrast was added as an input to the intensity model described above, it improved the prediction performance (fig. 4C), suggesting that these cells (at least partially) encode the contrast.

**Figure 4:**
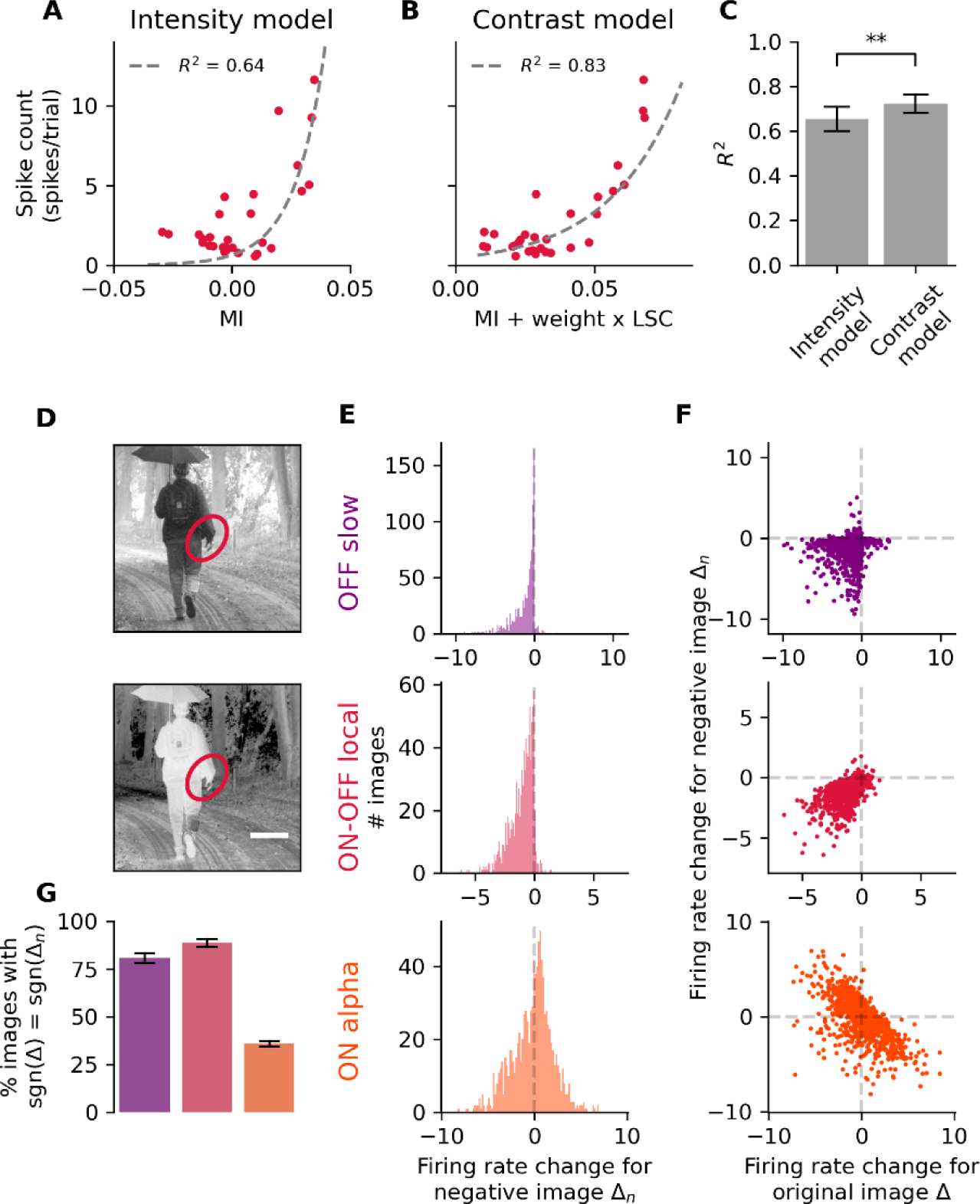
A simple contrast model and negative images confirm that defocus detectors encode contrast. **A**, Average firing rate as a function of mean intensity (MI) in the receptive field for an example ON-OFF local RGC (red dots) for the test set images (average firing rate over 30 presentations of the same image). The grey dotted line represents the fit of the intensity model. **B**, Same as panel A but for the mean intensity + local spatial contrast (LSC). **C**, distributions of the coefficient of determination R² for the intensity model (left bar, R² = 0.65 ± 0.05) and the spatial contrast model (right bar, R² = 0.72 ± 0.04) over all defocus detector cells (N = 15, p = 3 × 10^-3^, one-sided Wilcoxon signed-rank test). Data are represented as mean ± SEM. **D**, an example natural image (top) and its bright-dark inversed image (bottom). The ellipse represents the receptive field of an ON-OFF local cell (same cell as in panels A, B and E). **E**, distribution over 1000 images of the change of firing rate between 200 µm and –200 µm in response to the negative defocused images. Top, example OFF slow cell. Middle, example ON-OFF local cell. Bottom, example ON alpha cell. **F**, change of firing rate in response to the negative images vs. change of firing rate in response to the original images for the same cells as in panel E. Each dots represents an image. **G**, average (over cells) of the proportion of images leading to a firing rate change Δ_n_ (+200 µm vs. –200 µm), that as the same sign as the firing rate change Δ for the original image. Left, OFF slow (N = 4); middle, ON-OFF local (N = 3); right, ON alpha (N = 15). Data are represented as mean ± SEM. Data of panels **E, F** and **G** are shown for simulated central eye optics (0°). For simulated peripheral eye optics, see Supplementary figure 1.

To verify the hypothesis that defocus detector cells encode the contrast, we used an approach inspired from Goldin and colleagues (33). If a cell encodes contrast and not intensity, we expect a similar response to an image and its negative version (inverting bright and dark regions). Therefore, we used the CNN model to predict RGCs responses to the defocused versions of each natural image’s negative version. As observed for the set of original images, the firing rate was predicted to decrease when defocus switches from negative to positive for the two cell types we focused on (fig. 4E). This suggests that the responses of these cells to the defocus does not depend on the fact that the stimulus inside the receptive field is bright or dark. When plotting the change in firing rate triggered by a change in defocus in original and negative images, we found that these changes were highly correlated (fig 4F, G). This positive correlation between the response to an image and its negative strongly suggested that these cells encode spatial contrast rather than just luminance.

Note that, on the contrary, none of these properties were found for ON alpha cells. Firing rate could equally increase or decrease when defocus changed from negative to positive, as observed above. The responses to a normal image and its corresponding negative image were negatively correlated (fig 4F, G), as expected from a cell type where the response will scale with local mean intensity, rather than spatial contrast.

To summarize, these results show that defocus detectors RGCs compute the local spatial contrast in static natural images while this is not the case for all RGC types.

### Local spatial contrast allows defocus detectors to detect the sign of defocus

Since local spatial contrast seems to be necessary to understand the responses of the two cell types we characterized as defocus detectors, we tested if computing this spatial contrast is sufficient to estimate the sign of defocus. Starting with an example cell, we compare how the mean intensity and the LSC vary with defocus (fig. 5A). Depending on the image, the mean intensity can increase (red and green lines) or decreases (blue line) with defocus. It suggests that the average intensity inside the receptive field is not a reliable clue to detect the sign of defocus. On the contrary, for the three example images shown on fig. 5A, the LSC reaches a maximum for mild negative defocus and decreases with more positive defocus. It suggests that the LSC is a better clue to detect the sign of defocus, as its variation with defocus does not show dependence on the specific image.

**Figure 5:**
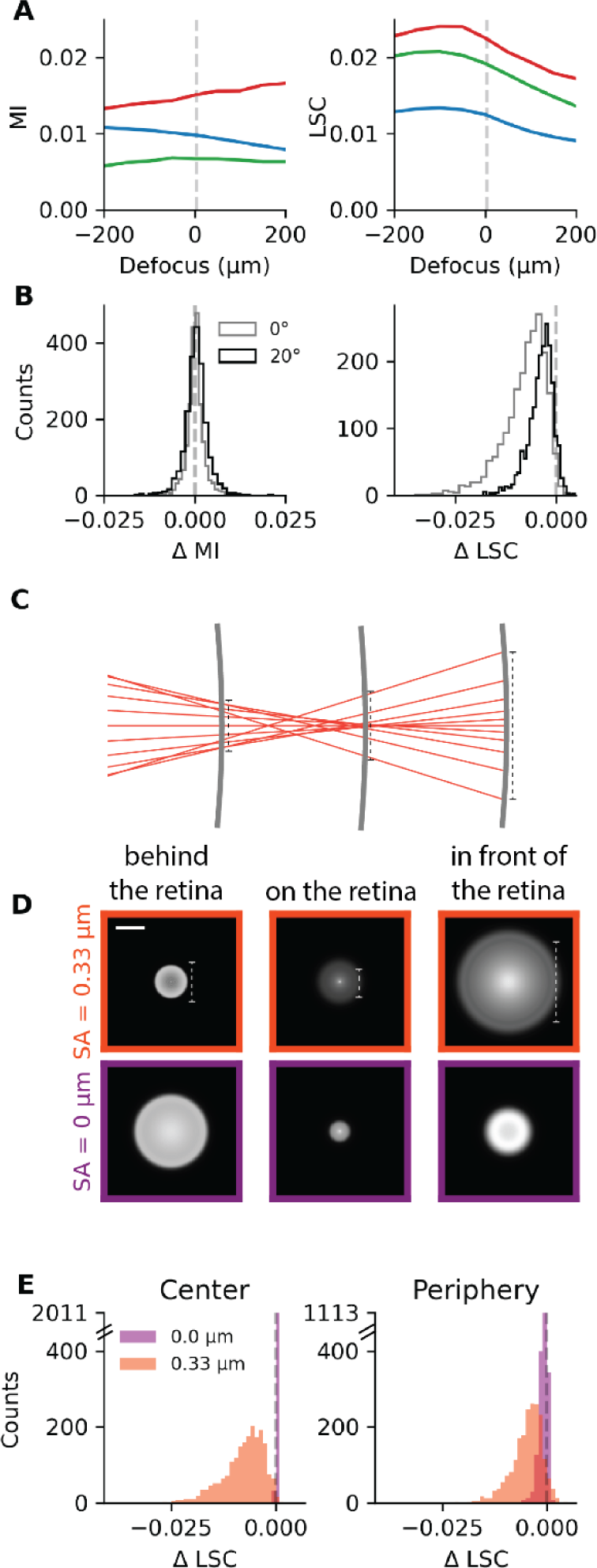
Spherical aberrations are necessary to detect the sign of defocus with the computation of local spatial contrast. **A**, mean intensity (MI, left) and local spatial contrast (LSC, right) as a function of defocus for three different images (red, green, blue curves, same color code as in figure 2C), computed in the receptive field of an example cell. **B**, distribution of the difference in mean intensity (left) and the difference in LSC (right) over N = 511 cells x 4 images, between defocus of −200 µm and 100 µm. Grey (black), in images corresponding to the simulated central (peripheral) eye optics. **C**, schematic illustrating the spherical aberrations in a mouse eye model with pupil diameter = 1.4 mm. The middle grey thick line represents a retina located at the focal point. The left (right) grey line represents a situation where the focal point would be located behind (in front of) the retina. **D**, top row, the PSFs with spherical aberrations (SA = 0.33 µm, PSF shown in log scale) corresponding to the positions of the three different retinas in panel C, for simulated central eye optics (0°). Bottom row, the PSFs at the same positions but without spherical aberrations. **E**, distribution of the difference in LSC over N = 511 cells x 4 images, with spherical aberrations (orange) and without (purple). Left (right), in images corresponding to the simulated central (peripheral) eye optics.

To confirm this observation at the population level, we measured the mean intensity and the LSC in defocused images in the receptive fields of many cells across 6 experiments (N = 511). To quantify the change of mean intensity and LSC across the cell population, we compute the difference in mean intensity (ΔMI) and LSC (ΔLSC) between 200 µm and −100 µm, around the maximum of LSC observed on panel 5A. We find that ΔMI can be positive or negative, depending on the cell, meaning that the mean intensity can increase or decrease with defocus (fig. 5B). It confirms that the mean intensity is not a reliable clue to detect the sign of defocus. On the contrary, ΔLSC is negative for almost all cells and images, meaning that the LSC almost always decreases with more positive defocus (fig. 5B). It confirms that the LSC is a reliable clue to detect the sign of defocus in natural images transformed by the eye optics and shows that computing the LSC allows to detect the sign of defocus.

Why is the LSC so different when the image is focused in front (positive defocus) or behind the retina (negative defocus)? We have shown above that the PSFs are very different in front and behind the retina (fig. 5D). These PSFs differ because the optics of the eye is not perfect. Of most importance are the spherical aberrations: all the rays that go through a lens do not focus at one single focal point (fig. 5C). This makes the blur asymmetrical with respect to the position of the retina. When the retina is in front of the focal point (left part of the schematic C), rays accumulate at the edge of the PSF, creating an intense ring at the border of the PSF (fig. 5D left). When the retina is located behind the focal point (right part of the schematic), rays at the edge of the PSFs spread on a larger area, so that the peak of the PSF is more intense that the edge, and the PSF is spread in a larger area (fig. 5D right). This difference allows a simple computation like spatial contrast to be sufficient to detect the sign of defocus. To test directly if spherical aberrations are important for defocus detection, we removed the spherical aberration from the simulated mouse eye model. This made the PSF in front or behind the focal point much more similar (fig. 5D). As a consequence, the LSC stopped decreasing systematically when switching from negative to positive defocus, and it became impossible to detect the sign of defocus by computing local spatial contrast (fig. 5E, see also supplementary fig. 2).

The spherical aberrations in the eye optics thus provide robust cues for defocus detection. To detect the sign of defocus, a realistic strategy for the retina could be to rely on spherical aberrations and on the computation of local spatial contrast.

### Local spatial contrast in human retinal images

Is the decrease in local spatial contrast when the focal point moves in front of the retina a general feature in realistic eye optics or is it specific to the mouse? To tackle this question, we developed a model of the adult human eye (see Methods) (fig. 6A). The point spread functions (PSFs) of the simulated central eye optics show the same features as in the mouse eye model: the PSF corresponding to the focal point being behind the retina is less spread than when the image is focused in front of the retina (fig. 6B). Consequently, when the image is focused in front of the retina (fig. 6B right), the LSC is smaller than when the image is focused behind the retina (fig. 6B left), for most of the tested natural images (fig. 6C, D) (see also (34)). We next tested whether the absence of spherical aberrations was also sufficient to cancel the change of LSC with defocus. If we remove spherical aberrations in the human eye model, the change in LSC is strongly reduced (fig. 6C, D). It suggests that computing the LSC could be a realistic strategy to detect the sign of defocus for the human retina too.

**Figure 6:**
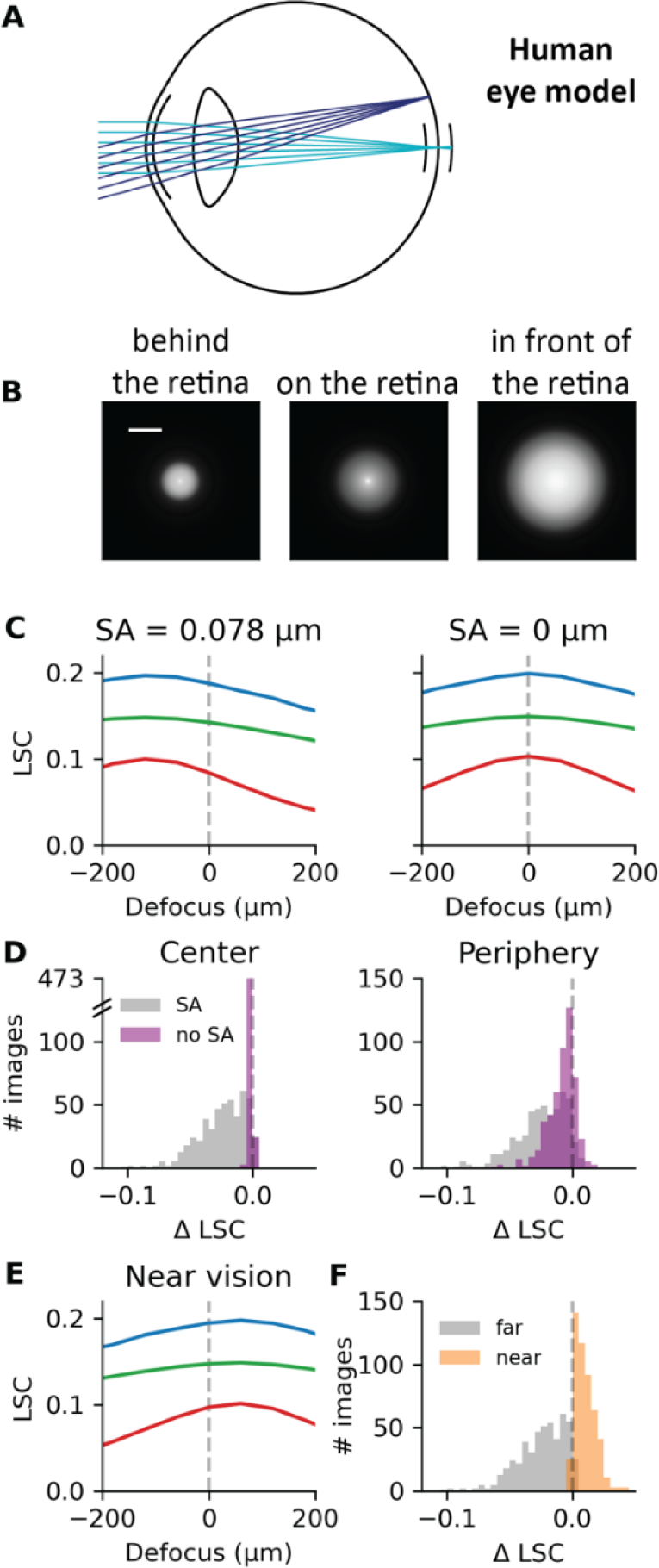
Near vision makes local spatial contrast more ambiguous to determine the sign of defocus. **A**, the human eye model. The light rays can reach the retina in the centre (0°, light blue) or periphery (20°, dark blue) of the retina. Pupil diameter is 5 mm. **B**, point spread functions of the simulated central eye optics when the image is focused behind (left), on (centre) or in front (right) of the retina. Scale bar, 50 µm. **C**, local spatial contrast (LSC) as a function of defocus for three example images (color code as figure 2C) with (left) and without (right) spherical aberrations, for simulated central eye optics (0°). **D**, distribution of the difference in LSC between 200 µm and −200 µm over N = 500 images, with spherical aberrations (grey, SA = 0.078 µm) and without (purple). Left (right), in images corresponding to the simulated central (peripheral) eye optics. **E**, LSC as a function of defocus for three example images (color code as figure 2C) in near vision (focus proximity = 2.5 D). **F,** distribution of the difference in LSC between 200 µm and −200 µm over N = 500 images, for a model with spherical aberrations and in far vision (grey, same as the grey histogram of panel D left) and a model with spherical aberrations in near vision (orange, focus proximity = 2.5 D, SA = 0.030 µm). In panel **E** and **F**, the LSC is calculated in images corresponding to simulated central eye optics (0°).

**Figure 7:**
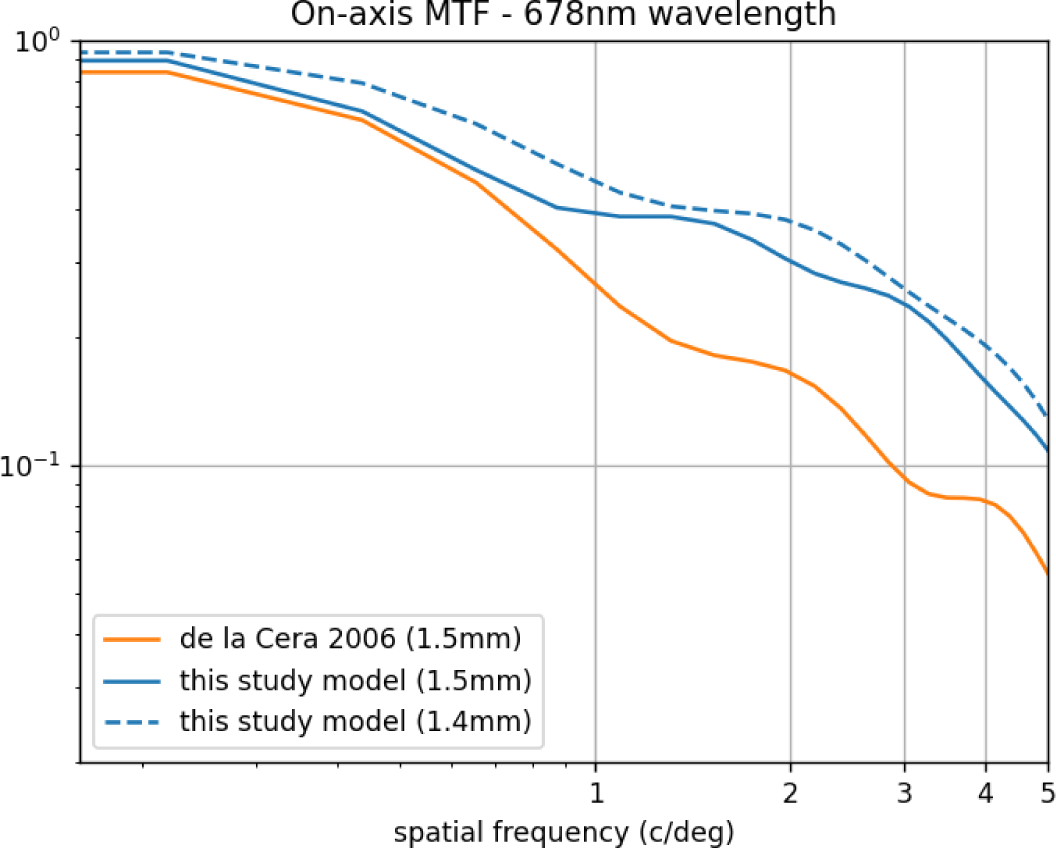
MTF radial profiles of the mouse eye model, simulated on-axis at a wavelength of 678 nm. Comparison with the results of Garcia de la Cera and colleagues (31) is made for a 1.5 mm pupil diameter, specifically from figure 4, line “mouse with defocus (1.5 mm)”. Moreover, the profile for a 1.4mm pupil is shown, which corresponds to the pupil size of interest in the current study.

In near vision, spherical aberrations are reduced, and can even change sign (35, 34). Therefore, we examined the effect of accommodation on the change in LSC with defocus. Simulating near vision in our human eye model, we found that the spherical aberration becomes slightly negative (−0.030 µm at 2.5 D of focus proximity vs. 0.078 µm at 0 D with a 5mm pupil) and that the LSC is higher when the image is focused in front of the retina (fig. 6E, F). If the retina relies on local spatial contrast to determine the sign of defocus, near vision will make this task more difficult.

## Discussion

We designed a new hybrid approach to record mouse retinal ganglion cell’s (RGCs) response to realistic defocused images with multi-electrode arrays (MEA). We simulated the optical transformation of the visual input using a detailed eye model that transformed the natural images that we presented to the retinas. We found that spherical aberrations of the eye optics caused an asymmetry between the image focused in front, and the image focused behind the retina. Due to these aberrations, ganglion cells that compute local spatial contrast could detect the sign of defocus because they systematically had a stronger response to a focus behind (negative defocus) compared to a focus in front (positive defocus). Estimating mean intensity alone was not sufficient to unambiguously estimate the sign of defocus. It is thus possible to estimate where the focal plane is by relying on two key features, spherical aberrations of the eye optics and local spatial contrast computation.

Taking into account the details of the transformation that the eye optics imposes on natural stimuli was necessary to understand how the retina could detect the sign of defocus. Previous works (36, 37) where the defocus was simulated using an artificial lens did not find a difference, or are inconclusive, probably because they did not have substantial spherical aberrations. As a result, the responses to negative and positive defocus could not be distinguishable (36). The detection of the sign of defocus relies on the specificity of the eye optics.

Our results suggest a possible algorithm for the retina to detect the sign of defocus, by relying on spherical aberrations and computing local spatial contrast: an overall decrease in local spatial contrast suggests that the focal plane is in front of the retina, while an increase suggest that the focal plane is behind it. We focused on two ganglion cell types whose response decreases when the focus went from negative to positive. We also found other types of ganglion cells that seem able to detect the sign of defocus by computing contrast (supplementary fig. 3), in particular ON-OFF ganglion cells. A previous study has shown that a specific ganglion cell type with a long latency seems to be suited for signaling whether an image is in focus or not (38), although it was thought that it could not make a difference between positive and negative defocus (39). In our study, each natural image was flashed for 300 ms, and this was too fast compared to the latency of this cell type. For this reason, we could not determine in our experiments if this cell type was also able to encode the sign of defocus. However, using stimuli having different temporal dynamics, and taking into account the spherical aberrations of the eye optics when presenting stimuli like we did in our study, it is possible that this type could also signal the sign of defocus.

The simplicity of this computation of local spatial contrast makes it probably widespread across cell types (see also (33)). While we have studied this computation in ganglion cells, local spatial contrast can also probably be computed by some amacrine cell types, especially the ones that receive both ON and OFF inputs (dopaminergic amacrine cells (40), vasoactive intestinal polypeptide-expressing amacrine cells (41, 42), nitric oxide synthase-containing amacrine cells (43), but also others). The modulation of eye growth by the retina does not seem to be caused by spiking ganglion cells (20, 21). For this reason, our results suggest that the retinal cell types that would be causally involved in modulating eye growth to achieve emmetropization could be amacrine cells that are able to compute local spatial contrast. Future works will be needed to determine which amacrine cell types can compute local spatial contrast.

While we don’t have causal evidence suggesting that our algorithm relying on spherical aberrations and local spatial contrast is actually the algorithm used by the retina to estimate where the focal plane is to modulate eye growth accordingly, our algorithm gives several predictions that have been verified by previous studies, and it also gives an explanation to several observations made in the literature about myopia. Myopia is an excess of eye growth, and an important public health concern. A major factor driving eye growth is the reduced illumination associated with indoor spaces (44). However, other factors are also associated with an excessive growth of the eye. Many studies have aimed at mitigating this health issue by slowing down eye growth (45). Among them, it has been proposed to have patients wearing glasses with a mild decrease of contrast in the periphery (46), a strategy that can slow down eye growth partially (47). If the retina relies on local spatial contrast to determine the sign of defocus, as we proposed, a mild decrease of contrast is equivalent to displacing the focal plane in front of the retina. Our algorithm predicts that this decrease in local spatial contrast should slow down eye growth, because it is equivalent to an image focused in front of the retina, which should trigger a reaction to “bring back” the focal plane to focus and thus achieve emmetropization. This is indeed observed in patients: wearing these glasses slows down the axial growth of the eye (47). In addition, optical analyses suggest that other types of lenses that are efficient at slowing down myopia progression in children also decrease the contrast on the retina (48). Our algorithm thus provides an explanation for the partial success of these therapeutic strategies: the retina could have its estimate of the focal plane biased to be more in front than it is and would therefore slow down the eye growth to compensate for this.

A factor that seems to increase the probability of being myopic is near sight vision (49), although it is always hard to separate from other factors like the level of illumination (44). Near sight vision reduces the amount of spherical aberration (35), and we have shown that this makes the determination of the sign of defocus more difficult. One possible explanation for the involvement of near sight as a risk factor for myopia might be due to reduced spherical aberrations, and thus the depth of focus will be more difficult to estimate. However, other factors that are correlated with near sight vision, like a decrease of level of illumination, are also associated with higher risks of myopia, and probably also play an important role. Conversely, orthokeratic treatments, which can mitigate strong myopia, increase spherical aberrations (50). Overall, while we cannot prove a direct causal link, our algorithm provides an explanatory basis for several studies aiming at slowing down eye growth to mitigate myopia.

One issue with our algorithm is that it seems at odds with the results obtained with Form Deprivation Myopia (FDM). Several studies have shown that a strong decrease in contrast, usually induced by placing a transducent diffuser in front of the eye (see (2) for a review), increases eye growth. Hence, it was hypothesized that increasing the contrast on the retina could inhibit eye growth (22, 1). However, these protocols of form deprivation usually induce a severe loss of contrast and acuity. How FDM depends on the level of attenuation of contrast and spatial frequency content is still unclear. It seems to depend on the exact protocol and on the species (4, 51–53). Compared to the studies where a mild decrease of contrast has been shown to slow down eye growth (see fig. 10A of (48)), Form Deprivation Myopia protocols induce a much stronger loss of contrast (see fig. 2 of (52)), which is far outside of the range of loss of contrast that can be induced by a change in focus as simulated in our experiments. It is thus plausible that FDM taps into a very different mechanism than the small loss of contrast that we simulated in our experiments and has thus consequences that are not related. This is further corroborated by several studies suggesting that the mechanisms of FDM and emmetropization are different (1).

Other features have been proposed to estimate the depth of focus. Chromatic aberrations could also be a clue because the focal plane for blue is in front of the focal plane for green and red. In principle, a comparison of blue spatial contrast versus red spatial contrast should allow determining where the focal plane is. Recent works suggest that modifying the chromatic content can help slow down eye growth in humans (54). At the cellular level, several types of cells in the retina have been shown to compare UV and green signals (55, 56), in particular in natural stimuli (57). A detailed simulation of chromatic aberration at various eccentricities remains to be done, but it might be possible to combine contrast cues and chromatic cues to refine even more the estimate of the sign of defocus. Following our “hybrid” approach, future works could simulate if chromatic aberrations in the periphery of the retina produce asymmetric PSFs, with a difference that can be detected by these cell types.

Our study was based on investigating how ganglion cells respond to flashed natural images. While this can approximate how ganglion cells will respond to stimuli right after a saccade (58–60), more complex types of temporal dynamics are present during natural visual experience. Specifically, between saccades, the visual input will be animated by fixational eye movements, which will strongly reshape the visual input (61). Other types of motion might also affect retinal processing. Future studies will have to understand whether these temporal dynamics, which are susceptible to trigger adaptation in retinal processing, can impact how the retina can extract the sign of defocus.

## Methods

### Mouse eye model

We based our optical mouse eye model on data from (62) limited to spherical dioptres. The model parameters, described in the table below, were fine-tuned to match the on-axis modulation transfer function (MTF) at very low spatial frequencies, as measured by (31). A reasonable match near 0.5 cycles/degree is important because it corresponds to the visual acuity of mice (63). Our model presents a primary spherical aberration of 0.42 µm for a 1.5 mm pupil at a 678 nm wavelength, as quantified by Zernike coefficient 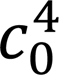, while (31) measured an average of 0.15 µm. A conic constant was later added as a parameter to the anterior cornea in order to investigate arbitrary spherical aberrations quantities, which allowed us to verify that similar effects were observable with lower amounts (see supplementary fig. 2). Finally, we introduced a dispersion formula for the crystalline lens, resulting in a longitudinal chromatic aberration (LCA) of 7.9 D between 457 nm and 633 nm, consistent with the findings of Geng and colleagues (30).

The PSFs are calculated using custom software. The simulation parameters include light source eccentricity, pupil size and defocus. To efficiently account for defocus, the image plane (i.e., the retina) is offset using the properties of the Fourier transform.

**Table 1:** Mouse eye model parameters.

| Component | Radius (mm) | Conic Constant | Distance (mm) | Refractive Index |
| --- | --- | --- | --- | --- |
| Anterior Cornea | 1.408 | $-1.38 + 3.15 * c_0^4$ | 0.092 | 1.402 |
| Posterior Cornea | 1.372 | 0 | 0.278 | 1.334 |
| Anterior Lens | 1.150 | 0 | 2.004 | $1.561 + 4.837e+03 / \lambda_{nm}^2$ |
| Posterior Lens | -1.134 | 0 | 0.942 | 1.333 |
| Retina | -1.598 | 0 | - | - |

### Human eye model

The human eye model is based on the model of Atchison (64), which has the specificity of having the ametropia as an input parameter. The conic constant of the anterior cornea was adjusted to match recent population spherical aberration data (mean 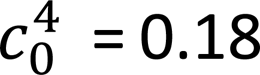 µm for a 6 mm pupil from (65) compared to 0.10 µm in the Atchison model). Moreover, the shape of the crystalline lens was also modified to correct the higher than observed peripheral astigmatism using data from (66). As for the mouse eye model, the light source eccentricity, the pupil diameter and de defocus can be varied. To simulate the accommodation process, the central curvature of the anterior lens was updated according to the stimulus proximity so that paraxial rays coming from an object point at near focus on the fovea.

### Polychromatic point spread functions

The eye models described above predict monochromatic PSFs at a given wavelength. As we wanted to simulate retinal images as seen by an eye watching at a natural scene, we computed more ecological PSFs using the sensitivity curves of the different types of cones, as follows. For the mouse, we considered only the M cone (peak intensity at 508 nm) and simulated PSFs for wavelengths ranging from 408 nm to 608 nm, every 10 nm. For the emmetropic human, we considered the M and the L cones (peak intensity at 537 and 567 µm, respectively) and simulated PSFs for wavelengths ranging from 450 nm to 650 nm, every 10 nm. To obtain the polychromatic PSFs, we then computed a weighted sum of the monochromatic PSFs, where the weight is the cones absorption value at the given wavelength.

### Electrophysiological recordings

All experiments were performed in accordance with institutional animal care standards of Sorbonne Université. We used six C57BL6J mouse retinas. After the euthanasia of the animal, the eye was enucleated and transferred into oxygenated Ringer medium (NaCl: 125 mol/L, KCl: 2.5 mol/L, MgCl2-6H2O: 1 mol/L, CaCl2: 1 mol/L, L-glutamine: 0.43 mol/L, NaH2PO4: 1.25 mol/L, glucose: 20 mol/L, NaHCO3: 26 mol/L) or Ames medium. Dissection was made into dim light condition. A piece of retina was mounted onto a dialysis membrane and then lowered with the ganglion cells towards the recording system. Retinas were kept at 36 degrees during the whole experiment.

Electrophysiological recordings of the ganglion cell’s spiking activity were subsequently performed with a multi-electrode array (252 electrodes, 30 µm distance between the electrodes, 8 µm diameter) as described in (33). In brief, the raw signal was acquired through MC_Rack Multi-Channel Systems software V 4.6.2 with a sampling rate of 20 kHz and high-pass filtered at 100 Hz. Spikes were isolated with SPYKING CIRCUS V1.0.9. Subsequent spikes analysis was performed with custom-made Python codes. We kept cells with less than 0.6% refractory period violation (2 ms refractory period). Cells that didn’t show a clear response to the white-noise stimuli (see below) were discarded because it prevents estimation of the receptive field and therefore the calculation of the local spatial contrast. In addition, we also discarded cells with a very low response to the chirp stimuli (see below) as it is needed for cell typing.

### Visual stimulations

A white mounted LED (MCWHLP1, Thorlabs Inc.) was used as a light source, and the stimuli were displayed using a Digital Mirror Device (DLP9500, Texas Instruments) and focused on the photoreceptors using standard optics and an inverted microscope (Nikon). The light level corresponded to photopic vision: 3.90*10³ and 1.48*10⁶ isomerisations/ (photoreceptor. s) for S cones and M cones respectively.

- Checkerboard stimuli

We displayed a random binary checkerboard during 40 min to 1 h at 30 Hz to map the receptive fields of ganglion cells, as described in (33). Check size was 42 µm and checkerboards had a size of 60 × 60 checks. Receptive fields were estimated using the classical ‘spike-triggered average’ (STA) method. A double Gaussian fit was performed on the resulting spatial STA, and the receptive field is defined as the ellipse corresponding to the 2σ contour of the fit.

- Natural image stimuli

We used the Open Access van Hateren Natural Image Dataset (67), which consists of 4212 monochromatic and calibrated images taken in various natural environments. Images were pre-processed as explained in (33).

The dataset of 3190 images remaining after the preprocessing step was used to stimulate the retina. We selected 30 images that were presented 30 times to create the test dataset for the CNN. The other images were only flashed once. Around 10% of them (250 images) were allocated to the validation dataset while the rest of the images composed the training dataset.

- Defocused images stimuli

To generate the defocused images, we selected 4 images in the above dataset and convolved them with the polychromatic point spread functions obtained with the mouse eye model described above. First, the image was upscaled to match the pixel size of the PSFs (1 µm). The upscaled image was convolved with the PSF using the fftconvolve function from the Scipy Python package. Then, the convolved image was downscaled to match the DMD pixel size (3.5 µm). Due to the position of the peak of the PSF that can change from one PSF to the other, some convolved images were shifted with respect to the original image. We re-aligned the convolved images on the original image to ensure that we recorded responses to a change of optical transformation and not to a shift of the image. Realignment was done by maximizing the correlation between the ‘contours Fourier image’ of the convolved image and the original. Briefly, Fourier transform was performed, low frequencies were filtered out and the inverse Fourier transform was applied, a similar process which is typically used to align calcium imaging data recorded with a 2-photon microscope. In some cases, small man-made adjustments were needed to ensure accurate realignment. Each defocused image was presented 25 times to the retina, randomly shuffled with the sharp natural images.

Images were displayed during 300 ms, followed by a grey frame presented during 300 ms. To obtain RGC’s response to the images, we counted the spikes occurring between 50 ms and 350 ms after the start of the image presentation.

### Cell typing

The cell typing procedure is described in detail in (33). In brief, in addition to the white-noise stimuli and the natural images, we displayed an additional full-field ‘chirp’ stimulus, similar to the one used by Baden and colleagues for their RGC classification (32). It is composed of ON and OFF steps of varying amplitude and frequencies with values ranging from 0 to 1. The chirp was presented 20 times during 32 seconds at a frequency of 50 Hz.

For typing the cells, we clustered the RGCs based on their response to the chirp stimuli and their STAs (33). It resulted in RGC groups that we assigned to one of the 32 types described in (32). To do so, we transformed our PSTHs in calcium events by convolving them with a decaying exponential (33; 68). Cell types that present strong responses to the modulating frequencies and amplitude were assigned correctly, while other types which mostly respond to ON/OFF steps, were assigned in a second round of correlation match. Besides the correlation of the chirp traces, we confirmed the correct assigning of groups by checking that the ellipses of each type form a proper mosaic, that the temporal STAs look uniform and the similarity of the spike waveforms. Finally, we correlated the chirp’s responses across experiments to ensure homogeneous groups.

### Ganglion cells modelling criteria

We modelled cells from retinas in which we found OFF slow and ON-OFF local cells (3 out of 6 retinas), and selected the cells following 2 criteria. First, having a proper STA. Second, having a stable response to the 30 sharp images that were presented 30 times randomly across the experiment. Stability was assessed using the criterion of (69). We computed the ratio between the explainable and the total variance and discarded all the cells with a ratio smaller than 0.22. In total, we were able to model 82 cells from 3 experiments.

### Convolutional neural network model

The architecture of the CNN model is described in detail in (33). Briefly, it is a two-layer network. The first layer is a convolutional layer made of four two-dimensional convolutional kernels. The filters are learned from the data independently for each experiment and common to all cells. The second layer is made of one filter for each cell, followed by a non-linearity.

The model was fitted on the data by minimizing a loss function on the training set, as explained in (33). The training set is made of 2910 sharp natural images and the corresponding RGC responses (number of spikes from 50 to 350 ms after the start of the presentation of the image).

To evaluate the performance of the model, we used a testing set composed of 30 sharp natural images that were presented 30 times and the associated RGC responses. Using repeated images allows to discriminate between the error due to the limitations in the model and the error due to the intrinsic noise in the RGC responses. We then calculated the noise-corrected correlation as introduced by (70) and detailed in (33).

### Classical LN model and SC model for mouse data

For this analysis, some additional cells were discarded because their STA was inaccurately fitted and to measure the local spatial contrast, receptive field estimation needs to be accurate. We consider the local image statistics, which are mean intensity and the contrast in the RGC’s receptive fields. Therefore, we consider only the pixels within the ellipse and transform the light level of each pixel in a Weber contrast: C = L-L_mean_/L_mean_ where L_mean_ is the mean light level over the entire image (28).

We used the classical LN model and the SC model developed by Liu and colleagues (28). We did exactly as they describe in their methods; therefore, we give only a brief description of the models here.

Both models start with spatial filters applied to the stimulus, F_LN_ and F_SC_, for the classical LN and the SC model, respectively. For each image, F_LN_ is the mean intensity in the receptive field, I_mean_.

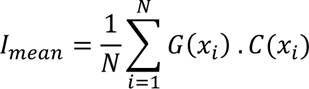

where the sum is taken over the N pixels positions x_i_ located within the receptive field (2σ contour of G(x_i_)) and C(x_i_) is the Weber contrast of pixel i. F_SC_ receives an additional input, the local spatial contrast (LSC), which is the standard deviation of the weighted pixels intensity in the receptive field:

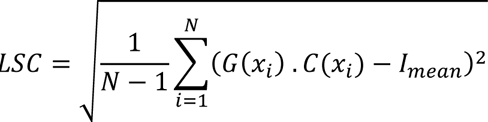

The LSC is added to I_mean_ with a weight w to obtain F_SC_

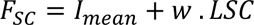

The firing rate was then predicted from F_LN_ and F_SC_ by passing them through a non-linear parametrized “softplus” function

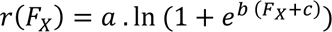

where F_X_ stands for F_LN_ or F_SC_. For each model, the responses to a training set of 3160 sharp natural images were used to fit r(F_X_). The parameters a, b and c were optimized with the curvefit function from Scipy with 50 iterations to find the initial values leading to the smallest residual error. The fitted functions were then used to predict the cell’s responses to the test set (30 images repeated 30 times). To quantify the model performance, we computed the coefficient of correlation R between the predicted response and the measured spike count and reported the explained variance R².

### Mean intensity and local spatial contrast in human retinal images

To obtain the LSC in natural images transformed with the eye optics of a human eye, we defined a generic receptive field as a 2D gaussian. The size of the receptive fields was chosen based on measurements in human retina (71). The receptive field of human parasol cells ranges roughly from 100 to 200 µm, with a median at 144 µm for ON cells and 123 µm for OFF cells. Human midget cells have a receptive field diameter ranging from roughly 30 to 100 µm, and the median for ON cells is 64 µm and 46 µm for OFF cells. We used 120 µm as standard size for 2-sigma contour of the gaussian fit. We then get the pixels comprised within that contour as an array of coordinates and compute the LSC with the above formula. We verified that, for the large majority of images, the method is robust to a variation of the size of the receptive field, within the range of the measured diameters.

## Acknowledgments

We would like to thank Matthew Chalks for his help with modelling and writing, and Ulisse Ferrari for his help on statistical analyses. This work was supported by ERC Consolidator grant DEEPRETINA (101045253), ANR grants (Chaire Industrielle MyopiaMaster ANR-22-CHIN-0006, ANR-18-CE37-0011 –DECORE, ANR-20-CE37-0018-04-Shooting Star, project NUTRIACT, project PerBaCo, project RetNet4EC), a grant from AVIESAN-UNADEV and one from Retina France to O.M. and by the ANR grant ANR-23-CE37-0004-01 HiDeepID to M.G. This work was supported by the Programme Investissements d’Avenir IHUFOReSIGHT 497 (ANR-18-IAHU-01).

## Author contributions

Conceptualization: SG, MG, KO, OM; Methodology: SG, SH, MG, KO, MO; Software: SG, SH, SV, MG; Formal analysis: SG; Investigation: SG, AW, TQ, MG; Visualization: SG; Supervision: KO, OM; Writing— original draft: SG, OM; Writing—review & editing: SG, OM.

