## Supplemental information for "Nonlinear spatial integration allows the retina to detect the sign of defocus in natural scenes"

Sarah Goethals *et al.*

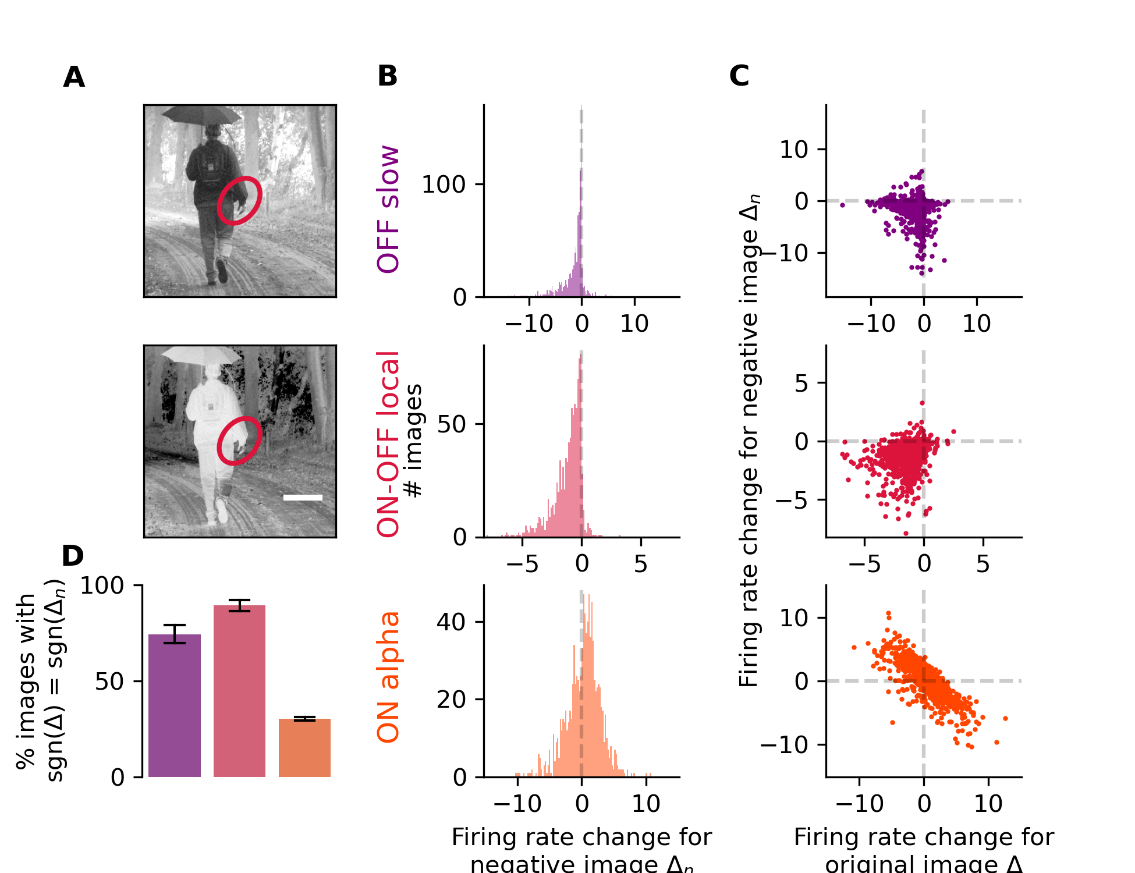

**Supplementary figure 1 -** A simple contrast model and negative images confirm that defocus detectors encode contrast, for simulated peripheral eye optics (20°). **A**, an example natural image (top) and its bright-dark inversed image (bottom). The ellipse represents the receptive field of an ON-OFF local example cell (same cell as in panels A, B and E). **B**, distribution over 1000 images of the change of firing rate between 200 µm and –200 µm in response to the negative defocused images. Top, example OFF slow cell. Middle, example ON-OFF local cell. Bottom, example ON alpha cell. **C**, change of firing rate in response to the negative images vs. change of firing rate in response to the original images for the same cells as in panel E. Each dot represents an image. **D**, average (over cells) of the proportion of images leading to a firing rate change ∆_n_ (between defocus of +200 µm and defocus of –200 µm), that as the same sign as the firing rate change ∆ for the original image. Left, OFF slow (N = 4); middle, ON-OFF local (N = 3); right, ON alpha (N = 15). Data are represented as mean ± SEM.

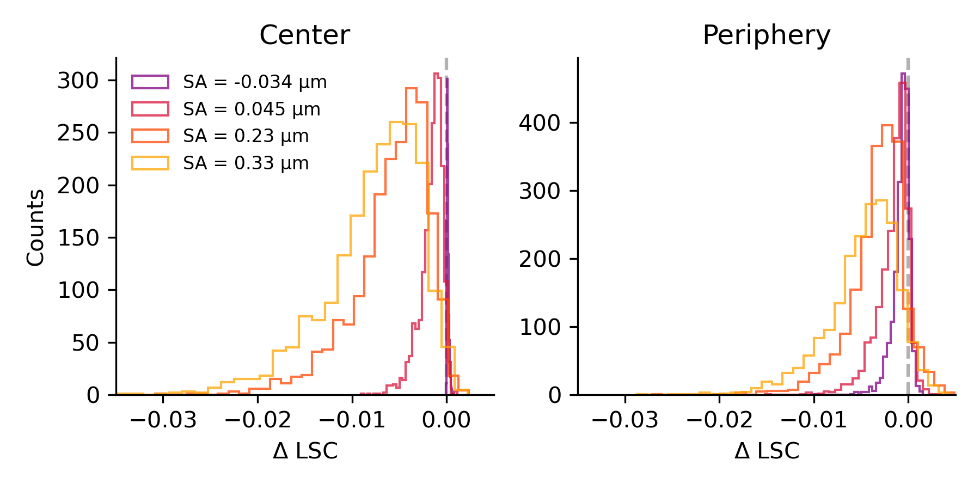

**Supplementary figure 2 -** Effect of variations of the amount of spherical aberrations in the mouse eye model on the local spatial contrast (LSC). Distribution over N = 511 cells x 4 images of the difference in LSC between a defocus of 200 µm and a defocus of -100 µm. Left, for simulated central eye optics (0°). Right, for simulated peripheral eye optics (20°).

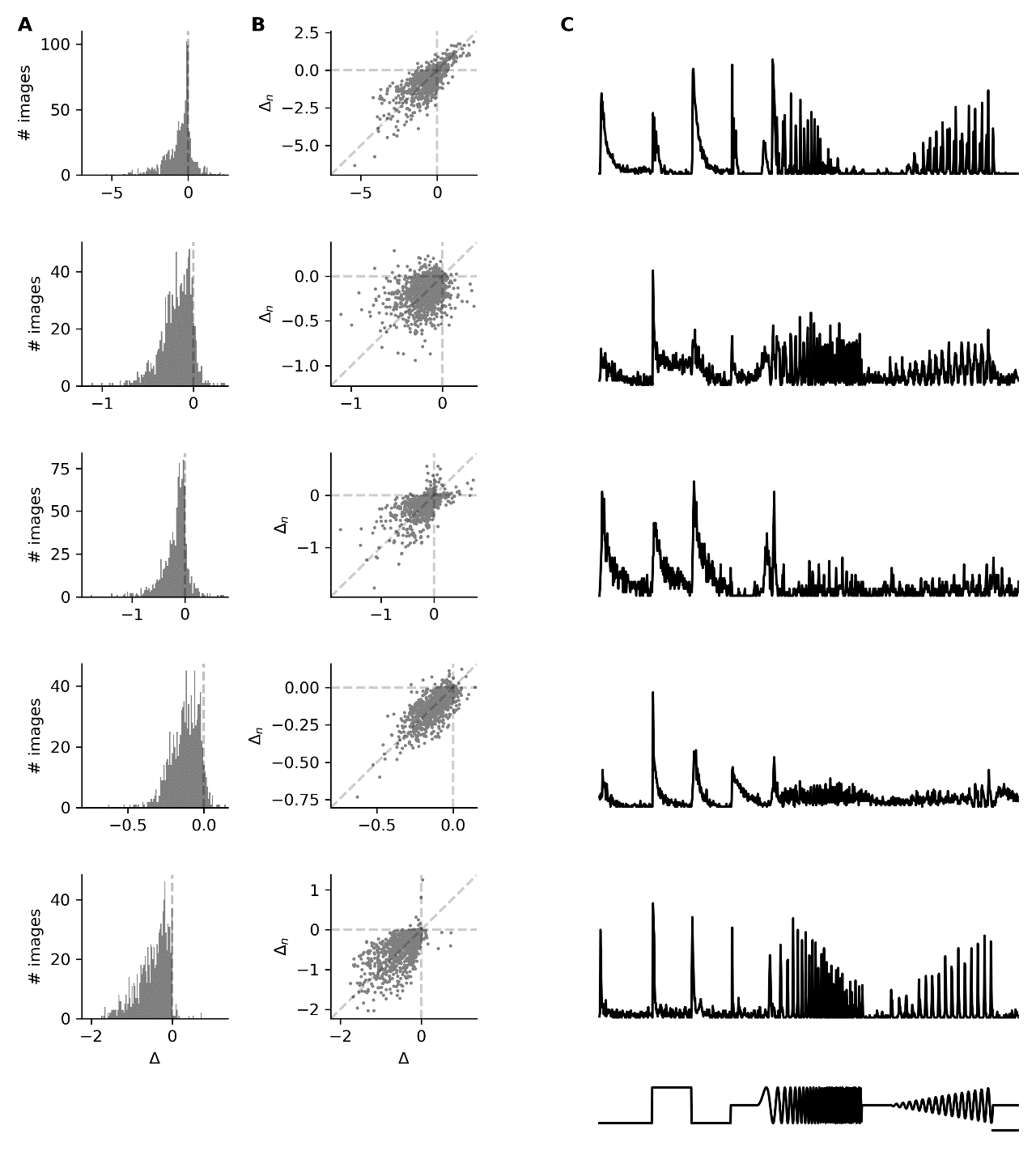

**Supplementary figure 3** - Defocus detectors can belong to other types. In addition to the OFF slow and ON-OFF local types in which all the cells are defocus detectors, we found defocus detector cells in other types. Each line corresponds to a different defocus detector cell. **A**, distribution over 1000 images of the difference in firing rate between positive (+200 µm) and negative (-200 µm) defocus. **B**, change of firing rate for the negative image vs. change of firing rate for the original image, for 1000 images. Each dot represents an image. **C**, response of the defocus detector cells to the bright-dark chirp (bottom trace). Scale bar: 2 seconds.
